# Practice Reshapes the Geometry and Dynamics of Task-tailored Representations

**DOI:** 10.1101/2024.09.12.612718

**Authors:** Atsushi Kikumoto, Kazuhisa Shibata, Takahiro Nishio, David Badre

## Abstract

Extensive practice makes task performance more efficient and precise, leading to automaticity. However, theories of automaticity differ on which levels of task representations (e.g., low-level features, stimulus-response mappings, or high-level conjunctive memories of individual events) change with practice, despite predicting the same pattern of improvement (e.g., power law of practice). To resolve this controversy, we built on recent theoretical advances in understanding computations through neural population dynamics. Specifically, we hypothesized that practice optimizes the neural representational geometry of task representations to minimally separate the highest-level task contingencies needed for successful performance. This involves efficiently reaching conjunctive neural states that integrate task-critical features nonlinearly while abstracting over non-critical dimensions. To test this hypothesis, human participants (n = 40) engaged in extensive practice of a simple, context-dependent action selection task over 3 days while recording EEG. During initial rapid improvement in task performance, representations of the highest-level, context-specific conjunctions of task-features were enhanced as a function of the number of successful episodes. Crucially, only enhancement of these conjunctive representations, and not lower-order representations, predicted the power-law improvement in performance. Simultaneously, over sessions, these conjunctive neural states became more stable earlier in time and more aligned, abstracting over redundant task features, which correlated with offline performance gain in reducing switch costs. Thus, practice optimizes the dynamic representational geometry as task-tailored neural states that minimally tesselate the task space, taming their high-dimensionality.

## Introduction

Practice enhances performance on a wide range of tasks, from motor skills to more cognitively demanding, goal-directed behavior (A. Newell, 1993; Shiffrin & Schneider, 1977), like formulating geometry proofs (Neves and Anders, 1980) or even writing books (Ohlsson, 1992). Understanding how expertise in control-demanding tasks is achieved is crucial for explaining cognitive control processes, as it constrains how we perform goal-directed behavior following a controlled-to-automatic continuum.

Performance improvements over the course of practice are widely assumed to reflect changes in the neural representations of a task and the computations that access and act on those representations. Specifically, improvement is often thought to result from strengthening associations among task-relevant information, while reducing the influence of task-irrelevant factors, thereby facilitating access and use of task-relevant response pathways (Cohen et al., 1990; Crossman, 1959; Mackay, 1982; Shiffrin & Schneider, 1977; Verguts & Notebaert, 2009). For example, the dominance of word reading, as seen in the Stroop effect, is commonly attributed to strengthened associations between written color words and their spoken labels, developed through extensive practice reading. In this way, reading becomes automatic when seeing a written word, allowing for highly efficient and accurate execution, even in the absence of attentional and control demands (Cohen et al., 1990; Moors & De Houwer, 2006; Stroop, 1935).

However, though many theories assume changes in task representations underlie practice, how and which representations change remains controversial. One reason for this fundamental gap in our understanding is that all tasks involve latent representations at multiple intermediate stages of computation, any or all of which could be the target of optimization (Du et al., 2022; Wood & Rünger, 2016; Zheng et al., 2024). Furthermore, the true objective functions of tasks are hard to infer even when the task structure is assumed to be fixed (Saxe et al., 2022). Accordingly, influential psychological theories of automaticity and skill learning, which comparably explain how cognitive control systems learn to reduce control and attentional demands while making performance more efficient and accurate, nevertheless differ substantively in terms of the level of task representations they assume change to improve performance.

For example, explanations for the power function-like improvement (i.e., rapid improvement in the early stage of learning with diminishing benefits over practice) in task performance have been one of the fundamental targets of theories of automaticity and skill learning (Gordon D. Logan, 1988; Allen Newell, 1990). Indeed, such a pattern of learning is so universally observed among tasks, that it is among the few contemporary laws cited in psychology (Gordon D. Logan, 1988; Teigen, 2002). Yet, theories accounting for this phenomenon differ in the levels of representation they assume change with practice.

Newell & Rosenbloom’s (1981) influential chunking theory proposed that practice yields chunking of task features at both intermediate and high-levels of complexity and specificity, which collectively leads to heightened efficiency. Thus, power law-like gains in performance follow from the early benefits of learning the simpler intermediate chunks that generalize to multiple task events, versus the complex chunks that are less frequently relevant. Other theories, similarly localize learning at an intermediate compositional level, and most commonly, in terms of context-independent stimulus-response (S-R) bindings (Schneider, 1985; Cohen, Dunbar, and McClelland, 1986).

In contrast, Logan’s (1988) instance theory of automaticity assumes improvement reflects a shift of behavioral control from a slow but flexible algorithmic process to a one-step direct retrieval of specific, high-level task-event instances that combine all task features, including the context. Practice adds instances to memory, which facilitates this retrieval process. In time, retrieval predominates, but with diminishing returns on faster performance. Thus, unlike the above cases, learning occurs only at the highest level (i.e., memory of individual events) and not at intermediate levels.

Thus, even for the most widely observed learning effects, multiple theories explain the same pattern of learning outcomes using different psychological constructs. Hence, a theoretically important question concerns the locus of representational change during practice that leads to behavioral improvement. What task representations are targeted by practice?

One approach to addressing this question is characterizing how practice optimizes the representational geometry of neural task representations and how these changes translate to improved performance (Johnston & Fusi, 2023; Li et al., 2024; Pang et al., 2023; Saxe et al., 2022). Theoretically, a geometric view of task representations could unify selective enhancement, reduction (or abstraction), optimization, and the underlying mechanisms of information coding and neural changes within a coherent framework (Badre et al., 2021; Duncker & Sahani, 2021; Farrell et al., 2022; Fusi et al., 2016; Kriegeskorte & Kievit, 2013; Musslick et al., 2020; Saxe et al., 2022). This perspective has the potential to bridge constructs in psychological theories (e.g., chunks and instances) to their neural computations. Yet, empirical evidence on characterizing how practice reshapes neural task representations over learning trajectories is still scarce (Mill & Cole, 2023; Wojcik et al., 2023).

Accumulating evidence suggests that the specificity and generalizability of task representations adapt to the task demands by shaping their geometry and dimensionality (Badre et al., 2021; Fusi et al., 2016; Jazayeri & Ostojic, 2021; Li et al., 2024), which are both important outcomes of practice. In general, high-dimensional task representations are associated with increased separability of task features and more efficient and accurate task performance. This is typically achieved by mixing separate coding axes for the combinations of task features in a structured manner (Bhandari et al., 2024; Fine et al., 2022; Ito et al., 2022; Kikumoto et al., in Press; Kikumoto & Mayr, 2020; Ritz & Shenhav, 2024; C. Tang et al., 2020; Weber et al., 2023), such as in a nonlinear conjunction of all task-relevant features (stimuli, responses, and contexts), that define a task event, or in a random and task-agnostic manner (Rigotti et al., 2013, 2010; E. Tang et al., 2019). In contrast, low-dimensional task representations typically benefit generalization or re-use of neural solutions. This is because task representations are abstracted over task-irrelevant features to generate compositional manifolds (Bernardi et al., 2020; Courellis et al., 2023; Driscoll et al., 2024; Flesch et al., 2022; Goudar et al., 2023; Hazy et al., 2007; Johnston & Fusi, 2023; Tafazoli et al., 2024; Vaidya & Badre, 2022) or compressed representations (Hummos et al., 2022; Muhle-Karbe et al., 2023).

What is unknown is how task representations evolve to these task-tailored neural states over the course of learning, particularly in tasks where nonlinear integrations of contexts with response pathways are required for successful performance (but see (Courellis et al., 2023; Mill & Cole, 2023; Wojcik et al., 2023). On one view, new tasks are initially computed compositionally, and over time, specific combinations are learned as context-specific, high-dimensional representations. To some extent, this change mirrors predictions by some theories of automaticity described above, in that progress to highly skilled behavior increasingly relies on a library of specific representations (e.g., chunks or instances). Conversely, however, early in task learning, expressive high-dimensional representations might make multiple readouts available for optimization (Farrell et al., 2023; Flesch et al., 2021). Over the course of practice, irrelevant subspaces could be compressed or aligned to avoid interference or redundancy (Cohen et al., 1990). This results in a transition toward a compositional and low dimensional geometry. As a third alternative, behavioral improvement may occur strictly at the handling of sensory and/or motor information or maintenance of contexts (Anderson, 1982; Cole et al., 2018; Meyer & Kieras, 1997; E. H. Schumacher, 2001).

Thus, there is a significant empirical gap in our understanding of how the representational geometry of task representations change in human brains throughout practice in ways that yield behavioral improvements. This issue remains unresolved because various theoretical constructs can equally well account for the same learning effects on behavior without clarifying underlying neural task representations. The current study addresses this gap in knowledge by directly relating changes in the geometry and dynamics of task representations to improvement in performance over three days of context-dependent action selection. Participants performed tasks requiring the nonlinear integration of rule context to select appropriate S-R (stimulus-response) mappings, shared across contexts (Mayr & Bryck, 2005). This setup allowed us to explore the learning of many task representations of varying complexity, including S-R mappings that are context-independent or context-specific (i.e., task-event file; Frings et al., 2020; Hommel, 2004; Eric H. Schumacher & Hazeltine, 2016; Verguts & Notebaert, 2009). Further, we mapped multiple task cues to the same rule context, which allowed us to test how such redundant task features become accommodated over practice.

We hypothesized that expertise in goal-directed actions arises from the optimization of the representational geometry in a task-tailored manner. Specifically, neural states maximally separating required context-specific task contingencies should be enhanced because they provide an efficient readout to solve the targeted task. Such neural states are equivalent to encoding context-specific conjunctions—nonlinear integration of task features such as the context and stimulus-response mapping to define a task event. In other words, practice should enhance stable readout of context-specific conjunctive neural states to improve task performance. Further, we hypothesized that these separable, conjunctive subspaces are gradually tailored to align unnecessary dimensions to abstract task features that are not critical to solving the task at hand, because such solutions are minimally high-dimensionality.

Consistent with these hypotheses, we found that practice optimizes the representational geometry of task representations selectively. Specifically, context-specific, high-level conjunctive representations were present early in practice and became stronger as a function of power-law, selectively accounting for rapid performance improvements, which was not the case for other, lower-levels of representation. Across days, these conjunctions recruited more separable yet stable dynamics at earlier points relative to the start of the trial. At the same time, these dynamics were reshaped to become more abstract across redundant task features, reducing the unnecessary dimensions of tasks. Overall, these findings suggest that practice optimizes encoding and/or retrieval of task-tailored, context-specific conjunctions. These conjunctions minimally orthogonalize contingencies to solve the task and slowly become abstract over redundant task features, serving as the locus of practice leading to expertise.

## Materials and Methods

### Participants

Forty participants (20 females; mean age: 23 years) were recruited and gave informed consent using procedures approved by the Human Subjects Committee at the RIKEN. All participants had normal or corrected-to-normal vision and had no history of neurological or psychiatric disorders. After preprocessing the EEG data, one participant was removed and was not analyzed further due to excessive artifacts (i.e., more than 25% of trials; see *EEG recordings and preprocessing* for details). Two participants did not complete the last session but their data for earlier sessions were retained for the analysis.

### Behavioral Procedure

The task closely followed the procedure in Kikumoto and Mayr (2020). Specifically, participants performed a cued rule-based action selection task (Figure 1A), which required participants to behave according to 16 unique rule-stimulus-response (S-R) conjunctions (Figure 1B). On a trial-by-trial basis, a randomly selected action rule (i.e., vertical, horizontal, clockwise, and counter-clockwise rule) specified four different S-R mappings, based on a simple spatial transformation over four possible stimuli (a dot at top-left, top-right, bottom-left, and bottom-right of the white frame). The spatial transformation indicated the target button-press response from four options that were arranged in 2 x 2 square matrix (4, 5, 1, and 2 on the number pad) that mirrored the stimulus display. Thus, for instance, the “vertical” rule mapped the top-left circle to the bottom-left response. Two different synonymous cue words were randomly used to specify action rules. Further, though the sets of four rule-stimulus-response conjunctions were unique for a given rule, individual S-R mappings could be shared across different rules. For example, just as with the vertical rule, the counter-clockwise rule would also map a circle in the top-left to the bottom-left response, but this rule would require a different S-R mapping than the vertical rule for a circle in the upper left or bottom right. Following the logic of Mayr and Bryck (2005), this design feature allowed us to distinguish the effects of practice on context-integrated conjunctive representations versus shared, compositional stimulus-response rules.

**Figure 1.**
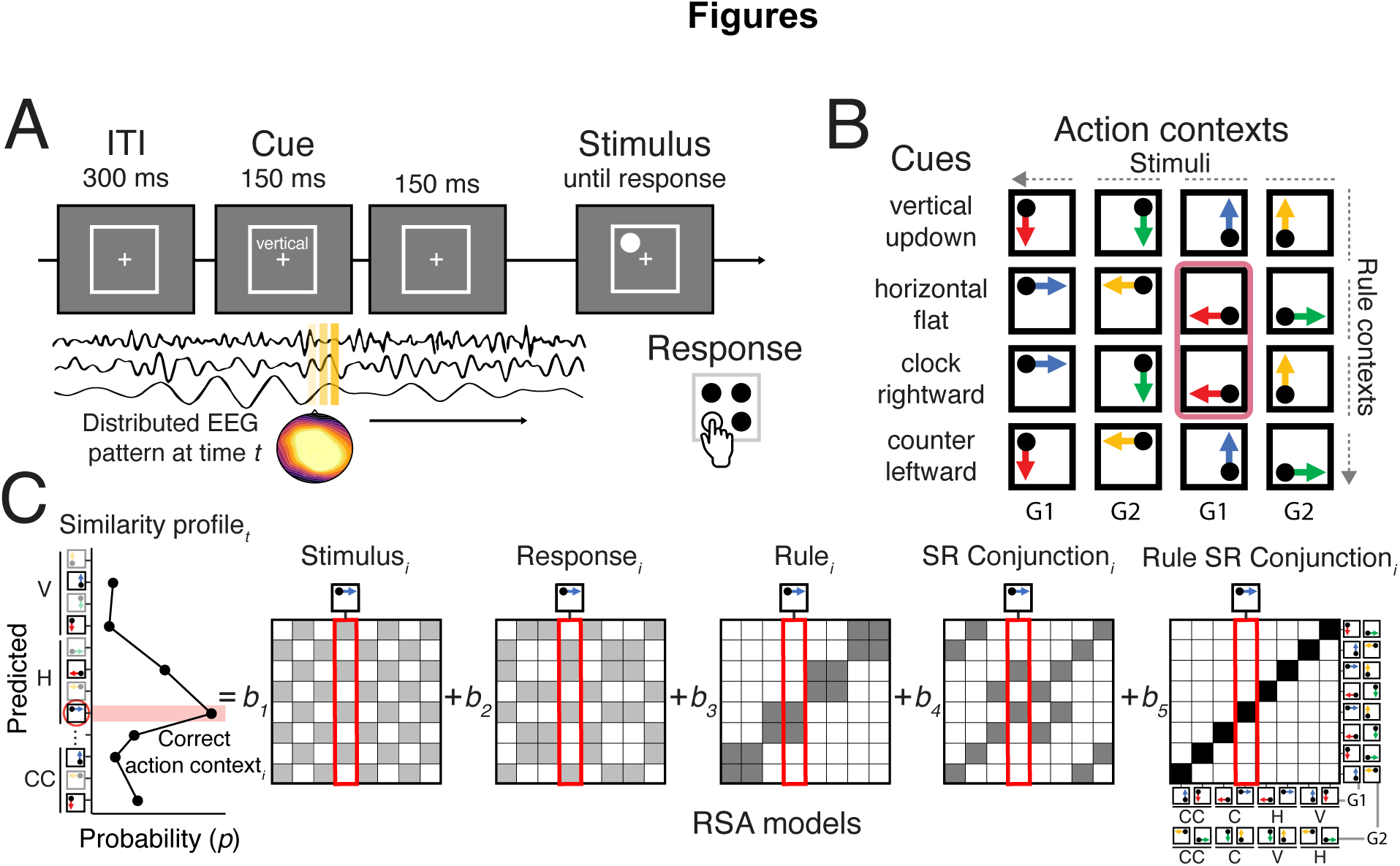
Task design and the procedure of decoding analysis. Both the task design and decoding analysis procedure is identical to Exp.2 in Kikumoto & Mayr (2020). (A) Sequence of trial events in the rule-selection task. (B) Spatial translation of different rules (rows) mapping different stimuli (columns) to responses (arrows), yielding 16 independent conjunctions. Two action contexts highlighted pink, for example, share the S-R mapping. (C) Schematic of the time-resolved representational similarity analysis (RSA). For each sample time (*t*), a scalp-distributed pattern of EEG was used to decode the specific rule/stimulus/response configuration of a required action. The decoder produced sets of classification probabilities for each of the possible action contexts. The profile of classification probabilities reflects the similarity structure of the underlying representations, where action contexts with shared features are more likely to be confused. For each trial and timepoint, the classification probabilities were regressed onto model vectors as predictors that reflect the different, possible representations. In each model matrix, the shading of squares indicates the theoretically predicted classification probabilities (darker shading means higher probabilities) in all possible pairs of action contexts. The coefficients associated with each predictor reflect the unique variance explained by each of the constituent features and two different types of conjunctions: stimulus-response (SR) conjunction and rule-specific SR (Rule SR) conjunction. To fully counterbalance the conditions, action contexts were grouped into two (G1 and G2), and then the RSA procedure was applied.

**Figure 2.**
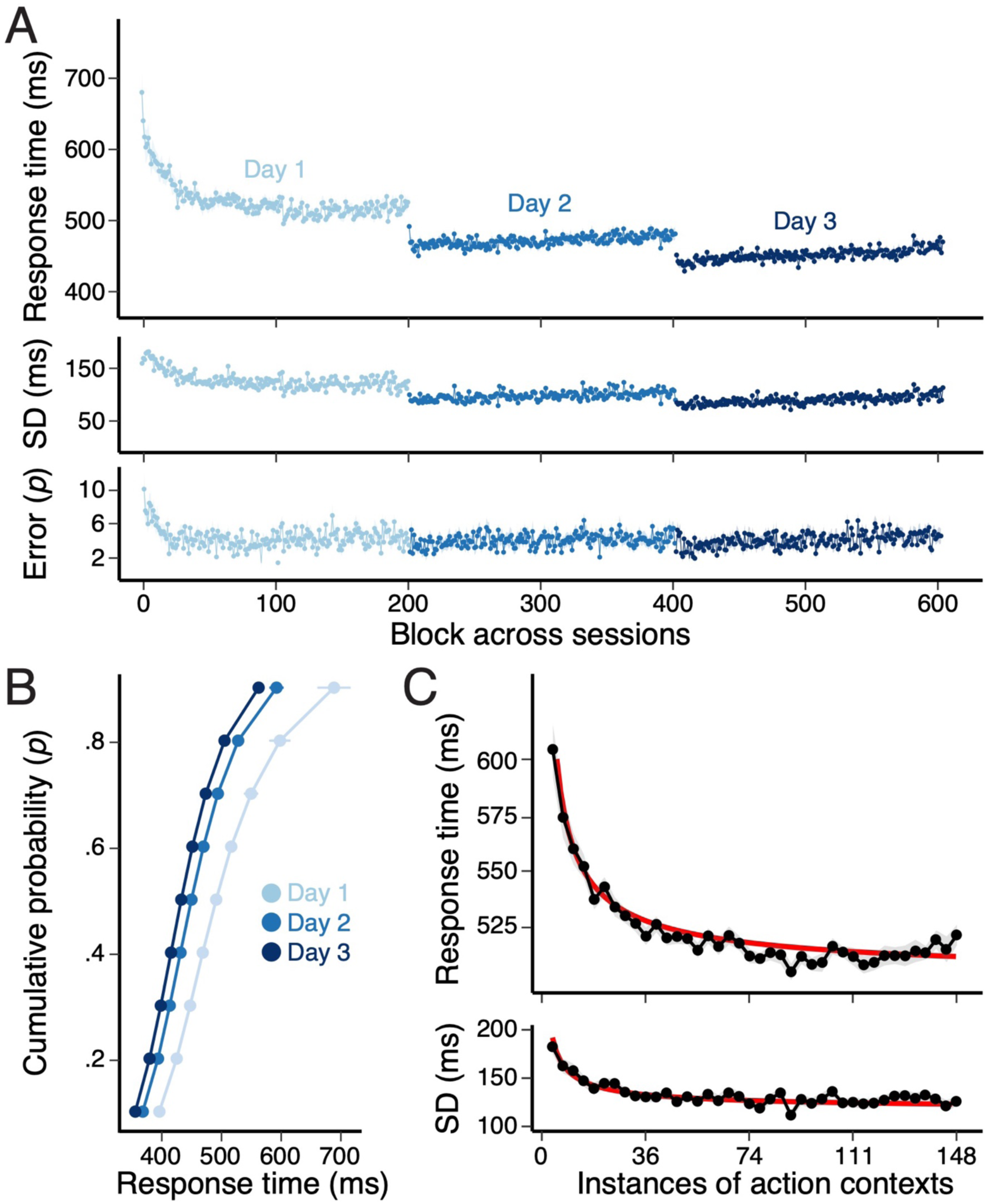
Practice effects on task performance. (A) The average response times (RTs), the variability (standard deviations or SDs) of RTs, and the probability of errors within each block. (B) Changes in RTs across days over the cumulative density of RT distributions on each day. (C) Changes in RTs and SDs on day 1 sorted by the binned instances of specific action contexts (Figure 1B). Each bin contains on average 40 trials. Red lines show the best-fitted power functions with the same exponent.

Participants were tested in three sessions across three consecutive days. Each session consisted of 202 experimental blocks, where the first two blocks did not provide performance-based incentives but were identical to other blocks. On the first day, participants practiced 5 blocks without EEG measurement. Participants were instructed to complete as many correct trials as possible that commenced within each 18-s block (trials that began within the 18-s window were allowed to finish). Participants were given a performance-based incentive for trials with RTs faster than the 75th percentile of correct responses in the preceding blocks when 1) the overall accuracy was above 90% and 2) there were more than 7 completed trials in a given block.

### EEG recordings and preprocessing

The continuous EEG was recorded using a Brain Products actiCHamp recording system (Brain Products GmbH). Recordings were obtained from a broad set of scalp sites (Fp1, Fp2, F7, F3, F4, F8, FC5, FC1, FC2, FC6, T7, C3, C4, T8, CP5, CP1, CP2, CP6, P7, P3, P4, P8, PO9, PO10, O1, O2, Fz, FCz, Cz, Pz, and Oz), from the left and right mastoids, and from electrodes lateral to the external canthi and below the left eye. The electrodes (TP9 and TP10) were placed on the mastoids beneath the cap as reference sites. Using additional passive electrodes, electrooculogram (EOG) was recorded from laterally placed electrodes to detect horizontal eye movements with the separate ground electrode placed on the left side of the forehead. The EEG was filtered online with .1 *Hz* high-pass and 1000 *Hz* low-pass filters and digitized at 1000 Hz. The left mastoid was used as a reference for all recording sites, and data were re-referenced off-line to the average of left and right mastoids.

The scalp EEG and EOG were amplified with an SA Instrumentation amplifier with a bandpass of .01–45 *Hz*, and signals were downsampled at 250 *Hz* using EEGLab (CITE). EEG data were first segmented into 22 s intervals to include all trials within a block. After time-frequency decomposition was performed, these epochs were further segmented into smaller epochs for each trial. For stimulus-aligned epochs, we used the time interval of -200 ms to 800 ms relative to the onset of stimuli. For response-aligned epochs, we used the time interval of -800 ms to 200 ms relative to the onset of responses. The trial-to-trial epochs including blinks (>250 *μ*v, window size = 200 ms, window step = 50 ms), large eye movements (>1°, window size = 200 ms, window step = 10 ms), blocking of signals (range = -.01 to .01 *μ*v, window size = 200 ms) were excluded from subsequent analyses. Any epochs that showed EEG activity greater than 5 s.d. were excluded.

### Time-Frequency Analysis

Temporal-spectral profiles of single-trial EEG data were computed via complex wavelet analysis (Cohen, 2014) by applying time-frequency analysis to preprocessed EEG data epoched for the entire block (>22 seconds to exclude the edge artifacts). The power spectrum was convolved with a series of complex Morlet wavelets), where *t* is time, *f* is frequency increased from 2 to 40 *Hz* in 35 logarithmically spaced steps, and σ defines the width of each frequency band, set according to *n*/2p*ft*, where *n* increased from 3 to 10. We used logarithmic scaling to keep the width across frequency band approximately equal, and the incremental number of wavelet cycles was used to balance temporal and frequency precision as a function of frequency of the wavelet. After convolution was performed in the frequency domain, we took an inverse of the Fourier transform, resulting in complex signals in the time-domain. A frequency band-specific estimate at each time point was defined as the squared magnitude of the convolved signal for instantaneous power.

### Representational Similarity Analysis

We performed a time-resolved multivariate pattern classification analysis to decode action-relevant information following our previously reported method with a few modifications (Kikumoto & Mayr, 2020). As the first step, at every sample in trial-to-trial epochs, separate linear decoders were trained to classify all possible action constellations (Figure 1C) for each S-R mapping. Specifically, we performed a penalized linear discriminant analysis using the caret package in R (Hastie et al., 1995; Kuhn, 2013). This step produced a graded profile of classification probabilities for each action constellation, reflecting the similarity of underlying multivariate neural patterns between actions.

Decoders were trained with the instantaneous power of rhythmic EEG activity, which was averaged within the predefined ranges of frequency values (2-3 *Hz* for the delta-band, 4-7 *Hz* for the theta-band, 8-12 *Hz* for the alpha-band, 13-30 *Hz* for the beta-band, 31-35 *Hz* for the gamma-band), corresponding to 155 features (5 frequency-bands X 31 electrodes) to learn.

Prior to the training of decoders, trials with incorrect responses were excluded. Within individuals and frequency-bands, the data points were *z*-transformed across electrodes to remove effects that scaled all electrodes uniformly. We used a k-fold repeated, cross-validation procedure to evaluate the decoding results (Mosteller & Tukey, 1968) by randomly partitioning single-trial EEG data into five independent folds with an equal number of observations of each action constellation. After all folds served as the test sets, each cross-validation cycle was repeated ten times, in which each step generated a new set of randomized folds. The number of observations in each target class (e.g., action constellation) was equated by randomly dropping excess trials for certain conditions. Resulting classification probabilities were averaged across all cross-validated results with the best-tuned hyperparameter to regularize the coefficients for the linear discriminant analysis.

We next performed representational similarity analysis (RSA) on the graded classification probabilities to assess the underlying similarity structure of task variables. The specific set of four rules ensured that each S-R mapping occurred in two different rules (e.g., a top-left circle leads to a bottom-left response in both the vertical and the clockwise rule), allowing us to decode context-specific SR conjunctions and rule-independent SR conjunctions separately (Figure 1B; Kikumoto & Mayr, 2022). Thus, each RSA model matrix represents the distinct similarity structure arising from different hypothetical, underlying representations (e.g., rules, stimuli, responses, SR conjunctions, and context-specific SR conjunctions; Figure 1C).

To obtain time-resolved estimates of each RSA model, we regressed the vector of logit-transformed classification probabilities onto RSA model vectors for independent decoding results. Importantly, to estimate the unique variance explained by competing models, we regressed all model vectors simultaneously. To fully counterbalance conditions and orthogonalize regressors, models were fitted within sub-group of action contexts (i.e., G1 and G2; Figure 1B), following the procedure in Kikumoto & Mayr, 2022. Further, individual time samples are concatenated within 12 ms non-overlatpping time windows to include subject-specific regressors of *z*-scored, average RTs, and error rates in each action context to reduce potential biases in decoding. These coefficients, expressed in their corresponding *t*-values, reflect the quality of action representations at the level of single trials, which was later related to variability in behavior and dimensionality measures. We excluded *t*-values that exceeded 5 SDs from the mean of each sample point. This procedure excluded less than 1% of all samples. These analysis steps were repeated separately using stimulus-aligned (Figure 3 and 4) and response-aligned EEG signals (Supplementary Figure 1).

**Figure 3.**
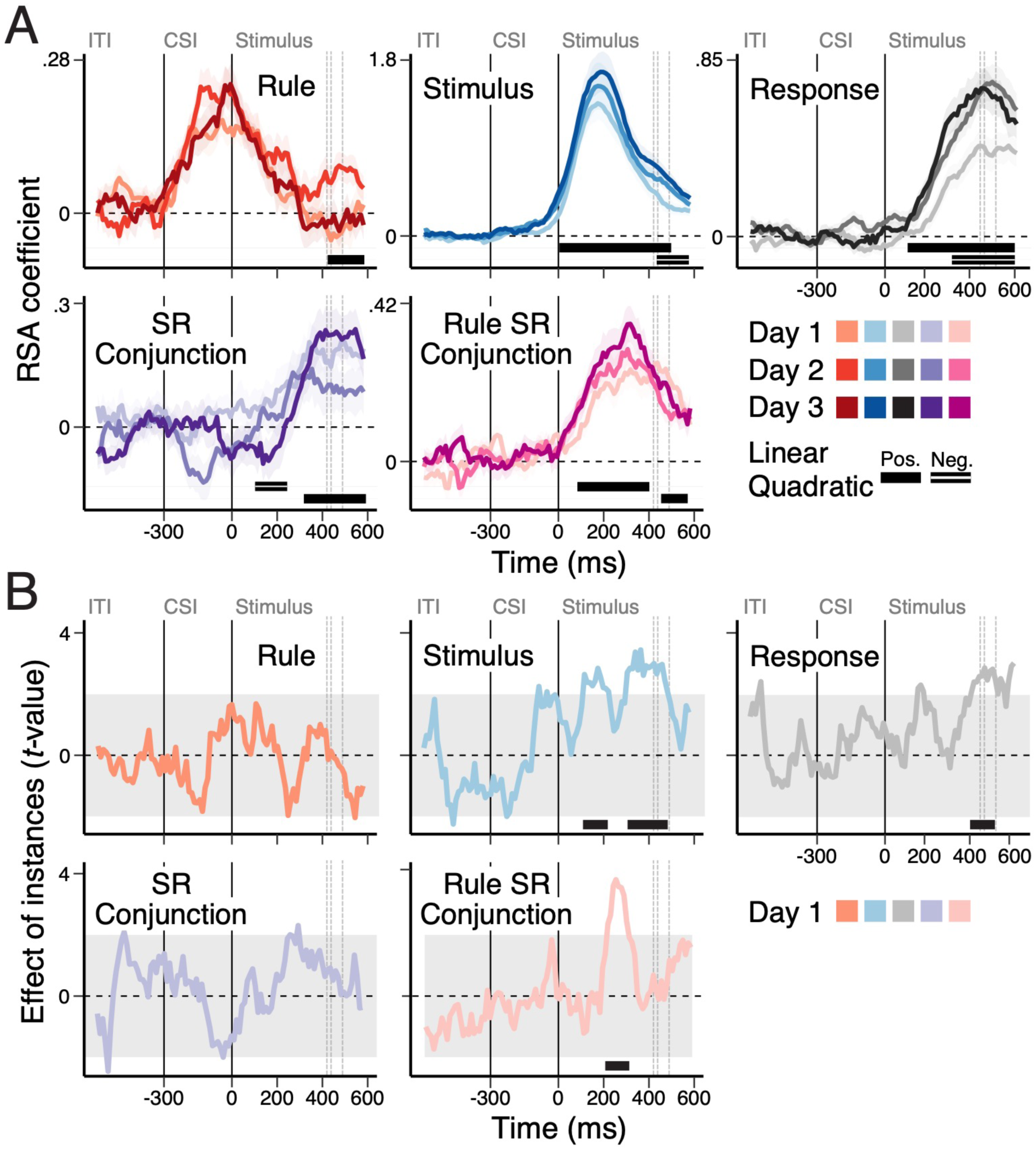
Stimulus-aligned time-course of decoding of task representations. (A) Average, single-trial RSA coefficients (*t*-values) associated with each of the basis set task features (rule, stimulus, and response) and two conjunctions (SR conjunction and rule-specific SR conjunction) that are aligned to the onset of the stimulus. Lines within each panel show the average RSA scores in each day. Shaded regions specify within-subject standard errors. (B) Modulation of RSA scores by the accumulated experiences of action context instances. (A-B) The black bars at the bottom of each panel denote the significant clusters using a non-parametric permutation test (cluster-forming threshold, *p* < .05, cluster-significance threshold, *p* < .01, two-tailed). Gray dotted lines show the mean RTs for each day.

**Figure 4.**
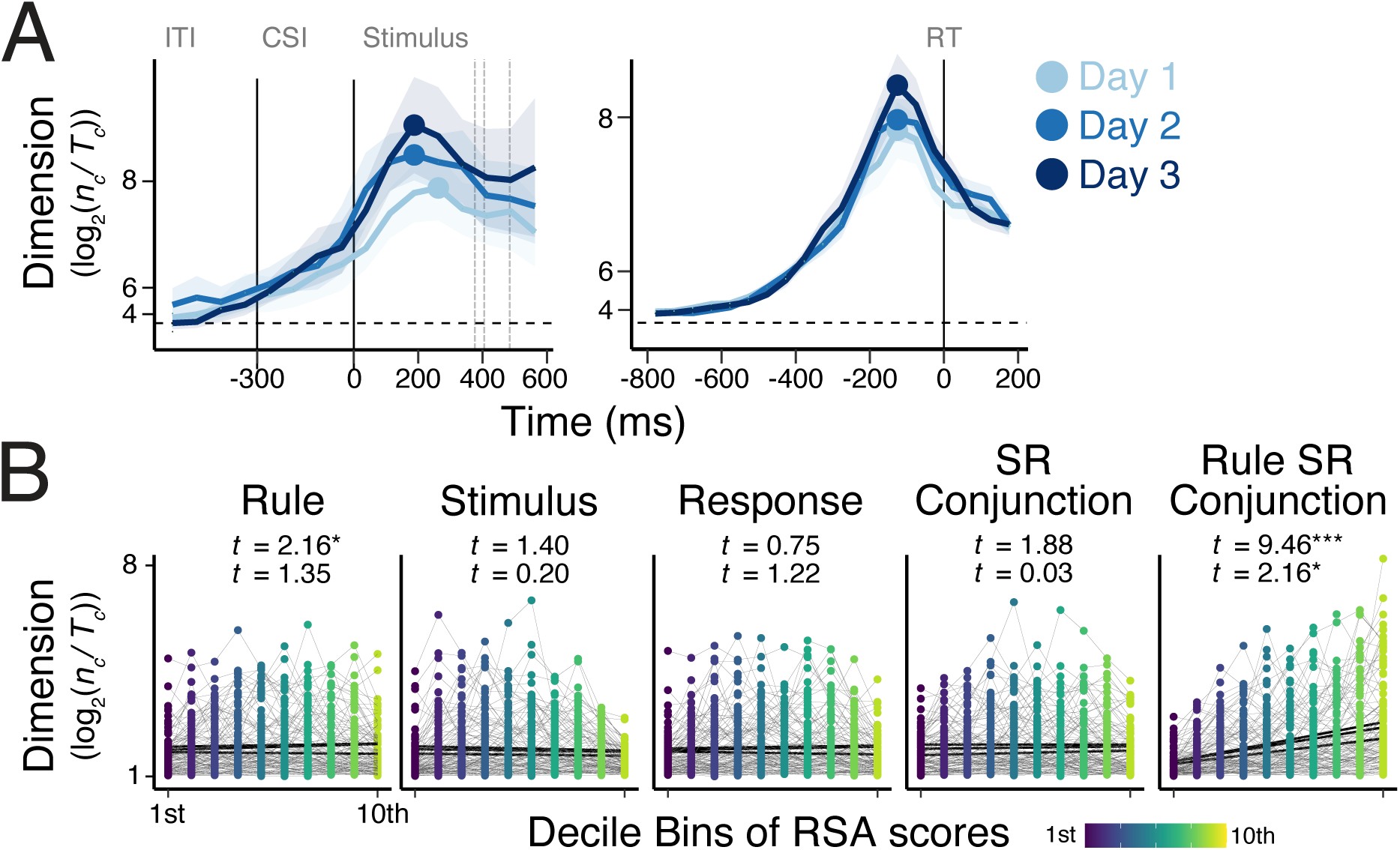
Task representational dimensionality. (A) Average, dimension score during action selection. The dimensionality is estimated as the proportion of implementable binary separation, *n_c_/T_c_*, out of all possible arbitrary binary pairs. The left and right panel correspond to the results using signals aligned to the onset of stimulus or responses respectively. Shaded regions specify within-subject standard errors. (B) Changes in the representational dimensionality as a function of RSA scores of each of the basis set task features (rule, stimulus, and response) and their conjunctions (SR conjunctions and rule SR conjunction). The dimension scores (*n_c_/T_c_*) are calculated separately in decile bins of RSA scores of each feature within subjects, which correspond to individual points in each panel. The two *t*-values within each panel are the statistic of a linear effect of an increase of RSA scores (top) and how it changes across days (bottom).

### Multilevel Modeling

We used multilevel models to further assess the decoded representations (Figure 3B; Table 1) and relate them to trial-to-trial variability in behavior. When dependent variables are sorted by the instances of specific action contexts or rule-stimulus-response conjunctions (e.g., to analyze the effects of learning on day 1), we used models with three levels of random effects: trials, different action contexts (i.e., individual context-specific SR mappings), and participants. When the effects of accumulated instances are irrelevant (i.e., the session effects), we dropped the random effects at the level of different action contexts. When RTs were used as the dependent variable, they were log-transformed. To assess the variability of RTs, the standard deviations within individual blocks (Figure 1A) or within bins of instances (Figure 1C) were used as dependent variables. For all models, random intercepts and slopes were included as random effects unless specified otherwise.

**Table 1.** MLM testing the effects of sessions (days) on task representations during early response selection (0-300 ms from the stimulus onset).

| Effects | Rule |  | Stimulus |  | Response |  | S-R Conjunction |  | Rule-specific Conjunction |  |
| --- | --- | --- | --- | --- | --- | --- | --- | --- | --- | --- |
|  | *beta* (CI) | *t*-value | *beta* (CI) | *t*-value | *beta* (CI) | *t*-value | *beta* (CI) | *t*-value | *beta* (CI) | *t*-value |
| Session (linear) | <.01 (-.01 .01) | .54 | .03 (.01 .06) | 2.70 | .02 (.01 .04) | 3.29 | -.02 (-.03 -.01) | -2.12 | .02 (.01 .04) | 2.34 |
| Session (quadratic) | <.01 (-.01 .01) | -.71 | <-.01 (-.02 .02) | -.08 | -.01 (-.02 .01) | -1.37 | -.01 (-.01 .01) | -.43 | -.01 (-.01 .01) | -.81 |

The RSA scores of action features were averaged over time intervals selected *a priori* based on previous studies: pre-cue period (-300 to 0ms from the onset of stimulus), early response selection period (0 to 300 ms relative to the onset of stimulus), and late response selection period (-300 to 0 ms relative to the execution of responses).

For the time-resolved regression (Figure 3B), we used non-parametric permutation tests to evaluate the decoding results in the time domain (Maris & Oostenveld, 2007). First, we performed a series of regression analyses over time and detected samples that exceeded the threshold for cluster identification (cluster-forming threshold, *p* < .05). Then, empirical cluster-level statistics were obtained by taking the sum of *t*-values in each identified cluster with consecutive time points. Finally, nonparametric statistical tests were performed by computing a cluster-level *p*-value (cluster-significance threshold, *p* < .05, two-tailed) from the distributions of cluster-level statistics, which were obtained by Monte Carlo iterations of regression analysis with shuffled condition labels.

### Temporal Generalization analysis

To assess the temporal stability of task representations, we used the temporal generalization analysis (Dehaene & King, 2016). First, we trained a set of decoders to classify action contexts, following the identical procedure as the time-resolved RSA analysis, using EEG patterns at the specific time point. These time-specific decoders were applied to the held out data obtained at identical (matched) or different (generalized) time intervals. This generates a series of decoding results in a two-dimensional matrix form where the on-diagonal entries correspond to the results in matched time points whereas the off-diagonal entries are the results of generalization of linear hyperplanes over other time. Assessing the pattern of temporal generalization over the course of learning reveals how the nature of underlying neural computation such as the temporal stability of linear readouts of task critical information changes. The resulting decoding results, corresponding to every pairings of training and testing non-overlapping time-intervals (window size = 8 ms), were submitted to the regression-based RSA procedure to decode rules, stimulus, response, SR conjunctions and context-specific conjunctions simultaneously. The results of stimulus-aligned and response-aligned data were generated in a similar manner as in the RSA steps.

### Dimensionality analysis via Binary Classification Method

To assess the representational dimensionality (i.e., shattering dimension) of task representations, we used a binary (pairwise) classification method (Rigotti et al., 2013; Bernardi et al., 2018), which evaluates the dimensionality by counting how many different input conditions (i.e., unique action contexts; Figure 1B) could be reliably separated or shattered into binary groupings by linear readouts (*n_c_/T_c_*). We adapted the original binary classification method introduced in Rigotti et al. (2013) to EEG data that contain a non-trivial level of noise by adding constraints on how we select implementable binary classifications (Kikumoto et al., 2023).

Specifically, we trained separate decoders for all possible 65536 (i.e., 2^16^ - 2, excluding two cases where all labels being identical) sets of arbitrary binary separations defined by 16 unique input conditions (Figure 1B). Resulting decoding accuracies were counted as reliable linear separations when the following two conditions were satisfied: 1) the decoding accuracies exceeded the arbitrary chosen cutoff threshold of decoding accuracy of .51 where the chance level is .50 and 2) above-threshold decoding accuracies were observed in each of the unique input conditions exclusively. This second criterion was applied to increase the sensitivity to detect changes in dimensionality scores driven by the non-linear mixing of information even when the cutoff threshold accuracy is significantly lowered to the chance level. Note that the effect of different cutoff method was tested previously ((Kikumoto et al., 2023), which did not affect differences in dimensionality across compared conditions in empirical data.

As a result, we obtained the proportion of reliable binary separations out of all possible binary groupings (*n_c_/T_c_* = count of implementable binary classifications, which approximates the dimensionality of neural responses to the input space (dimension = log2(*n_c_/T_c_*)). When comparing across days, the aforementioned exclusive cutoff was applied separately. The other settings of decoding analyses such as the steps of cross-validation and the exclusion criteria of trials were identical to the RSA analysis except for that EEG signals were averaged over 50 ms non-overlapping consecutive time windows to reduce the number of interactions.

## Results

### Practice effects on behavior

Within the first session, the average response time (RT), error rates, and variability of RT (i.e., standard deviations of RT in each experimental block) decreased over experimental blocks and approached an asymptote, following a power-like function. These changes were reliably explained by both linear and quadratic effects (Figure 2A; Mean RT: *t*(1,38) = -5.57, *beta* = -.41, 95 % CI [-.56 -.28] for linear effect, and *t*(1,38) = 7.02, *beta* = .32, 95 % CI [.23 .40] for quadratic effect; Error rate: *t*(1,38) = -1.16, *beta* = -.31, 95 % CI [-.79 .20] for linear effect, and *t*(1,38) = 4.80, *beta* = .91, 95 % CI [.54 1.28] for quadratic effect; Variability of RT: *t*(1,38) = -3.64, *beta* = -.93.16, 95 % CI [-1.43 -.43] for linear effect, and *t*(1,38) = 5.63, *beta* = .84, 95 % CI [.54 1.13] for quadratic effect).

To investigate learning trajectories within each action context, we sorted trials into 40 bins by counting cumulative encounters (i.e., instances) of correctly executed responses in action contexts corresponding to the occurrence of a specific rule-stimulus-response conjunction. Both RT and the variability of RT decreased as the number of experienced instances of specific conjunctions increased (Figure 2C; Mean RT *t*(1,38) = -4.21, *beta* = -.21, 95 % CI [-.30 -.11] for linear effect, and *t*(1,38) = 6.62, *beta* = .17, 95 % CI [.12 .22] for quadratic effect; Variability of RT: *t*(1,38) = -1.92, *beta* = -.26.16, 95 % CI [-.53 .01] for linear effect, and *t*(1,38) = 4.62, *beta* = .31, 95 % CI [.18 .44] for quadratic effect). These results indicate that task performance is rapidly optimized within the first session among independent action contexts in accord with the expected power-law pattern.

Comparing the first, second, and third days, both the average RT and variability of RT substantially decreased whereas error rates remained near floor (Figure 2A; Mean RT: *t*(1,38) = -6.40, *beta* = -.15, 95 % CI [-.20 -.10] for linear effect, Error rate: *t*(1,38) = 7.28, *beta* = -.24, 95 % CI [-.30 -.17] for quadratic effect; *t*(1,38) = -1.84, *beta* = -.02, 95 % CI [-.05 .01] for linear effect, Variability of RT: and *t*(1,38) = -.72, *beta* = -.01, 95 % CI [-.05 .02] for quadratic effect for error rates; *t*(1,38) = -6.40, *beta* = -.15, 95 % CI [-.20 -.10] for linear effect and *t*(1,38) = -7.28, *beta* = -.24, 95 % CI [-.30 -.17] for quadratic effect). The linear improvements in RTs were observed in all steps of vincentized RT distributions (Figure 2B). Though, within the later sessions, performance became significantly worse as the experimental session progressed (*t*(1,38) > 4.43), possibly due to fatigue. Nevertheless, these results indicate that after initial rapid improvement, task performance gradually improved over sessions. Because such learning effects are apparent at the beginning of later sessions, this improvement was due to task offline gains between sessions.

Next, we assessed the patterns of repetition priming costs in switching rule, stimulus, and response feature dimensions across trials. A lack of repetition benefits under switching of any task features (i.e., partial overlap of task features) across trials has been considered evidence of task feature binding (i.e., event-files: (Frings et al., 2020; Hommel, 2004; Mayr & Bryck, 2005; Rangel et al., 2023) and has been linked to the formation of conjunctions integrating goals or contexts (Kikumoto & Mayr, 2020). Consistent with these studies, we found that switching in any task feature dimension incurred substantial costs on behavior compared to trials when the rule, stimulus and response repeats completely, (Supplementary Figure 1; Mean RT: *t*(1,38) = 15.16, *beta* = .39, 95 % CI [.33 .44] and Error: *t*(1,38) = 13.43, *beta* = .87, 95 % CI [.7 1.0]). These effects persisted across sessions without substantial changes, Mean RT: *t*(1,38) = - 1.64, *beta* = .04, 95 % CI [-.09 .06] for linear effect; *t*(1,38) = 1.48, *beta* = .03, 95 % CI [-.01 .06] for quadratic effect, and Error: *t*(1,38) = -1.99, *beta* = .87, 95 % CI [.7 1.0] for linear effect; *t*(1,38) = 1.40, *beta* = .13, 95 % CI [-.05 .30] for quadratic effect. A direct comparison of complete repetition trials and all-switching trials also did not significantly change across days (*t*(1,38) < 1.50).

However, although the overall benefits of stimulus-response (S-R) repetitions compared to switching did not change significantly across days (*t*(1,38) < .16), the cost on RT of switching rules in S-R repetitions trials was significantly reduced across days, Mean RT: *t*(1,38) = -4.12, *beta* = -.03, 95 % CI [-.05 -.01] for linear effect; *t*(1,38) = 2.71, *beta* = .03, 95 % CI [.01 .04] for quadratic effect, and Error: *t*(1,38) = .97, *beta* = .07, 95 % CI [-.06 .20] for linear effect; *t*(1,38) = -.64, *beta* = -.05, 95 % CI [-.02 .08] for quadratic effect. These results suggest that learning across days reduced the costs of switching action rules most substantially.

Finally, we tested how trial-to-trial changes in action-relevant, but redundant features, affected performance. Specifically, two different cues signaled the same task rule. Thus, while using the cue to identify the task was important, differentiating between these cues was not critical to solving the task. Nevertheless, consistent with past observations of cue encoding effects (Logan and Arrington), when the task cue changed from one trial to the next, but all other task features (i.e., the rule, stimulus, and response) remained identical, costs in RTs, errors, and variability of RTs were evident (Figure 1C; Mean RT: *t*(1,38) = 11.18, *beta* = .67, 95 % CI [.55 .79]; Error: *t*(1,38) = 7.15, *beta* = .77, 95 % CI [.56 .99]; Variability of RT, *t*(1,38) = 11.18, *beta* = .67, 95 % CI [.56 .79]). Such cue-dependent switch costs were significantly reduced across days for the mean RTs and variability of RTs (Mean RT: *t*(1,38) = -3.43, *beta* = -.36, 95 % CI [-.57 -.15] for linear effect, and *t*(1,38) = .71, *beta* = .07, 95 % CI [-.13 .29] for quadratic effect; Error: *t*(1,38) = -1.47, *beta* = -.28, 95 % CI [-.65 .09] for linear effect, and *t*(1,38) = 1.81, *beta* = .34, 95 % CI [-.03 .70] for quadratic effect; Variability of RT: *t*(1,38) = - 3.48, *beta* = -.36, 95 % CI [-.57 -.15] for linear effect, and *t*(1,38) = .71, *beta* = .07, 95 % CI [-.13 .28] for quadratic effect). Thus, aligned with the improvement across days in general task performance and switching of action rules, action selection became more resilient to differences in task features that are redundant to the objective of the task.

### Task representations at an early stage of learning (day 1)

As session one (Day 1) showed strong power law-like improvement within the session, we first focused on analysis within this session. Over the course of action selection, we characterized how task representations of task rules, stimuli, responses, stimulus-response (SR) conjunctions, and context-specific SR conjunctions (i.e., Rule SR conjunction) change along with the improvement of task performance via single-trial RSA (Figure 1C).

Temporal trajectories of task representations closely followed the pattern observed previously (Kikumoto & Mayr, 2020; Figure 3A). During the cue-to-stimulus interval, cued rules were encoded. Then, during action selection after the stimulus onset, context-specific conjunctive subspaces emerged that are dissociable from other lower-dimensional task representations. All of these events preceded response execution (Supplementary Figure 2), replicating the previous studies (Kikumoto & Mayr, 2020; Kikumoto, Bhandari, Shibata & Badre, in press).

Next, we directly tested how accumulated instances of specific action contexts changed the strength of encoded task representations. We found that as the function of the logarithmic increase of the number of instances or episodes of successful context-specific actions, the strength of context-specific SR conjunctions increased at the early stage of action selection (i.e., 200 - 300 ms after the onset of stimulus; Figure 3B). This effect on context-specific conjunctions remained significant after controlling for trial-to-trial RT as a nuisance predictor, *t*(1,38) = 2.38, *beta* = .02, 95 % CI [.01 .05]. A similar effect of the number of instances was observed for encoding stimulus and response representations in the late response selection period (Figure 3B); however, such effects were not observed for SR conjunctions.

We further found that enhancement of context-specific conjunctions had a significant effect on how rapidly performance improvement progressed on Day 1. Specifically, stronger context-specific conjunctions were associated with fewer instances-to-asymptote in mean RT, *t*(1,38) = 2.19, *beta* = .01, 95 % CI [.01 .07] for the interaction effect, though not in the variability of RT, *t*(1,38) = .65, *beta* = <.01, 95 % CI [-.02 .03]. No such relationship was significant for other task representations. Thus, these results indicate that encoding context-specific conjunctions is enhanced by accumulated experiences in specific action contexts on Day 1, and this accumulation is directly related to classical power-law improvements in performance within the early session.

### Changes in task representations across days

Next, we considered how representations changed over the course of sessions (Days 1, 2, and 3). Across days, we observed that task representations at specific stages of action selection were selectively enhanced (Figure 3A). During the cue-stimulus interval, the strength of rule representations remained the same across days, *t*(1,38) = .18, *beta* < .01, 95 % CI [-.01 .01] for linear effect and *t*(1,38) = -1.54, *beta* = -.01, 95 % CI [-.01 .01] for quadratic effect. In contrast, during an early stage of response selection (0 to 300 ms after the stimulus onset), rule SR conjunctions were significantly enhanced, along with stimulus and response representations (Table 1). Importantly, these changes over days were absent for rule representations, and the strength of stimulus-response (S-R) conjunctions significantly decreased across days within this period (Figure 3A). However, this decrease in SR conjunctions was driven by shorter durations of activation from the stimulus onset, and the strength remained similar at the time responses were executed (Supplementary Figure 2). In fact, their strength increased briefly after response execution. These results indicate that each task representation is optimized at variable rates of change while participants gain expertise.

Although most task representations, in particular context-specific SR conjunctions, were consistently negatively correlated with trial-to-trial RTs across all sessions (Supplementary Figure 3), we did not observe significant interactions between behavioral changes across sessions and the strength of moment-by-moment RSA scores, (Rule SR conjunction: *t*(1,38) = -.84, *beta* = -.01, 95 % CI [.04 .02] for linear effect; *t*(1,38) = -.84, *beta* = -.01, 95 % CI [.04 .02] for quadratic effect; SR conjunction: *t*(1,38) = -.89, *beta* = -.06, 95 % CI [-.05 .02] for linear effect; *t*(1,38) = -1.22, *beta* = -.02, 95 % CI [-.05 .01] for quadratic effect). Further exploratory analyses also confirmed that the effects are absent even when focused only on the early phase of each session where the influence of fatigue-like patterns in later sessions is minimal (Table 2). Thus, unlike the within-session changes in performance at was tied to the accumulation of instances, we did not find evidence that offline gains are due to factors affecting decodability of task representations.

**Table 2.**
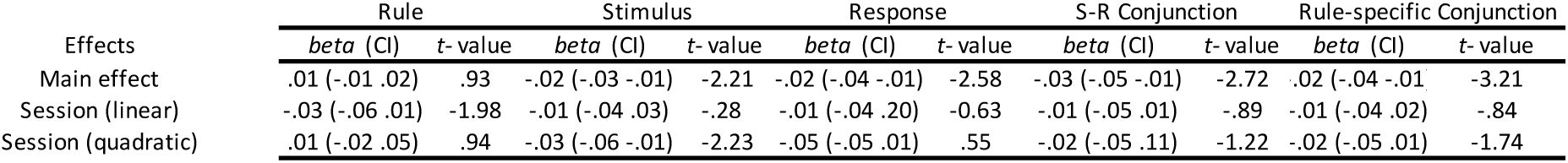
MLM testing the effects of task representations (0-300 ms from stimulus onsets) on RTs across sessions, focusing on early phase (first 75 blocks).

### Changes in representational dimensionality across days

Further, we assessed how practice changes the overall representational dimensionality of the task input space (i.e., unique action contexts) across days and how their changes relate to the quality of individual task representations. We previously observed that the higher dimensional formats are associated with faster and accurate responses, which peaked around 100 ms before response execution (Kikumoto et al., 2023). Across days, a similar temporal trajectories were replicated; however, the dimensionality only marginally increased at the peak across sessions (Figure 4A; *t*(1,38) = 1.45, *beta* = .18, 95 % CI [-.06 .41] for the linear effect; *t*(1,38) =-.08, *beta* = .01, 95 % CI [-.25 .28] for the quadratic effect). This increase correlated with encoding of stronger context-specific conjunctions (Figure 4B). Because both 1) increase of signal to noise ratio in encoding of context-specific conjunctions and 2) addition of diverse mixing of task features (i.e., heterogenous mixing) expand the representational dimensionality, these results suggest that practice did not diversity the mixing of task features rather enhanced context-specific conjunctions (but see **Changes in task-redundant dimensions**).

### Changes in representational dynamics across days

We previously observed that reaching stable neural dynamics in conjunctive subspaces is an important precondition for efficient context-dependent action selection (Kikumoto, Bhandari, Shibata & Badre, in press). If practice leads to better access to task memories providing direct solutions, the neural dynamics leading to orthogonal subspaces of goal states of the task (i.e., context-specific conjunctions) should be reshaped to be more efficient to provide stable readout for outputs. Specifically, the stable subspaces should be established earlier during action selection, and the subsequent readout should be resilient to task features that are not informative to define required goal states.

Consistent with this hypothesis, we found that the temporal stability of context-specific conjunctions increased across days (Figure 5). From Day 1 to Day 2, the stability generally increased during early response selection (0 ms to 300 ms after the stimulus onset). From Day 2 to Day 3, the forward generalization (i.e., the upper-left portion of the generalization matrix) is further enhanced, suggesting that the neural patterns during action selection become more stable at an earlier timing relative to the onset of stimulus (Table 3).

**Figure 5.**
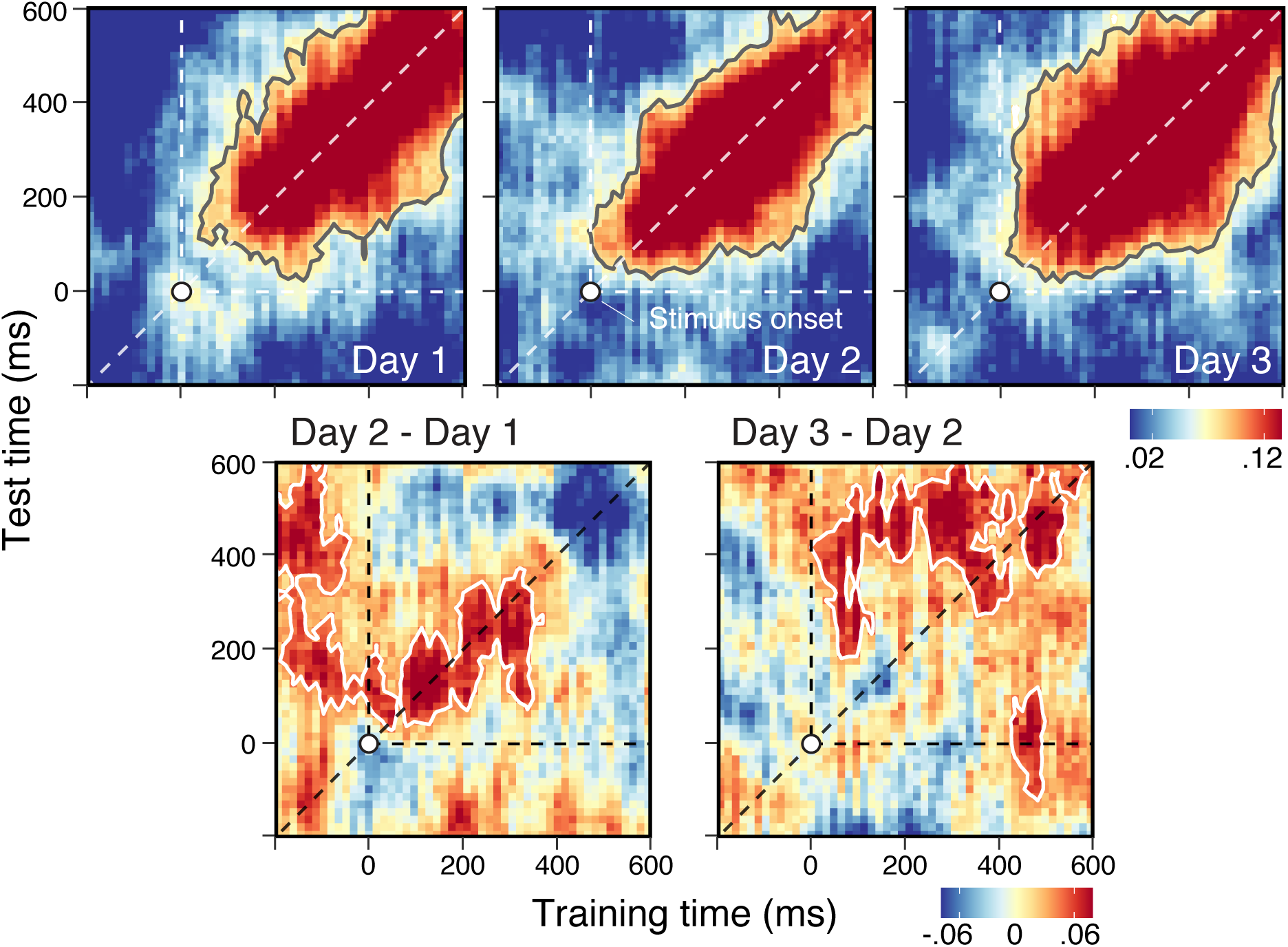
Representational dynamics in the subspace of conjunctive control representations. Temporal generalization matrix of linear hyperplanes that separate rule-specific conjunctions. The X-axis denotes time points where signals were sampled to train decoders then submitted to single-trial RSA. The Y-axis denotes time points where the test sets were derived. The regions surrounded by a gray/white contour denote the significant clusters using a nonparametric permutation test (cluster-forming threshold, *p* < .05, cluster-significance threshold, *p* < .01, two-tailed). The top panels show changes across days and the bottom panels show the differences between consecutive days.

**Table 3.**
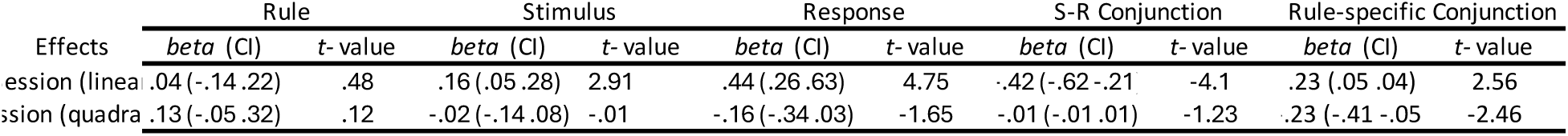
MLM testing the effects of sessions (days) on cross-cue generalization (0-300 ms from the stimulus onset).

More stable conjunctive subspaces were correlated with faster trial-to-trial RTs, *t*(1,38) = -2.85, *beta* = -.02, 95 % CI [-.28 -.01], controlling for the effect of time-resolved strength of conjunctions, *t*(1,38) = -3.40, *beta* = -.02, 95 % CI [-.36 -.01] or other task representations, replicating prior observations (Kikumoto, Bhandari, Shibata & Badre, in press). Further, the stable component showed a marginally significant interaction correlating with changes in RT across sessions, *t*(1,38) = -1.82, *beta* = -.02, 95 % CI [-.38 .01], such that more evidence of stable dynamics predicting better improvement across days. In contrast, the dynamic component showed a marginally significant interaction in the opposite direction, *t*(1,38) = 2.05, *beta* = -.02, 95 % CI [.09 .04], meaning that more evidence of dynamic representations predicted less improvement across days, however this difference between the stable and dynamic component on RTs across sessions was not statistically significant, *t*(1,38) = 1.49, *beta* = .018, 95 % CI [-.01 .04]. Thus, these results indicate that the neural dynamics that reach stable conjunctive subspaces are enhanced by extensive practice, which leads to better task performance.

### Changes over task-redundant dimensions

Finally, we tested whether optimal task representations are formed not only by separating context-specific conjunctive subspaces but also by compressing redundant dimensions within rule contexts. Specifically, we tested the degree of abstraction by characterizing alignment across task-redundant cues (Figure 1B).

As noted above, switching task cues incurs significant costs, even when the rule context and stimulus-response mapping stays identical (Jost et al., 2013; Mayr, 2006; Meiran, 1996). These costs were reduced over practice (Figure 6B; see ***Practice effects on behavior***). We further found that, across days, the context-specific conjunctions become more generalizable (i.e., abstract) across cues in both the dynamic and stable components of the dynamics (Figure 6B), *t*(1,38) = 2.56, *beta* = .23, 95 % CI [.05 .41] for linear effect and *t*(1,38) = -2.46, *beta* = -.22, 95 % CI [-.41 -.05] for quadratic effect (see Table 3 for other task features), suggesting practice over days reorganized the dynamics to be similar across task cues.

**Figure 6.**
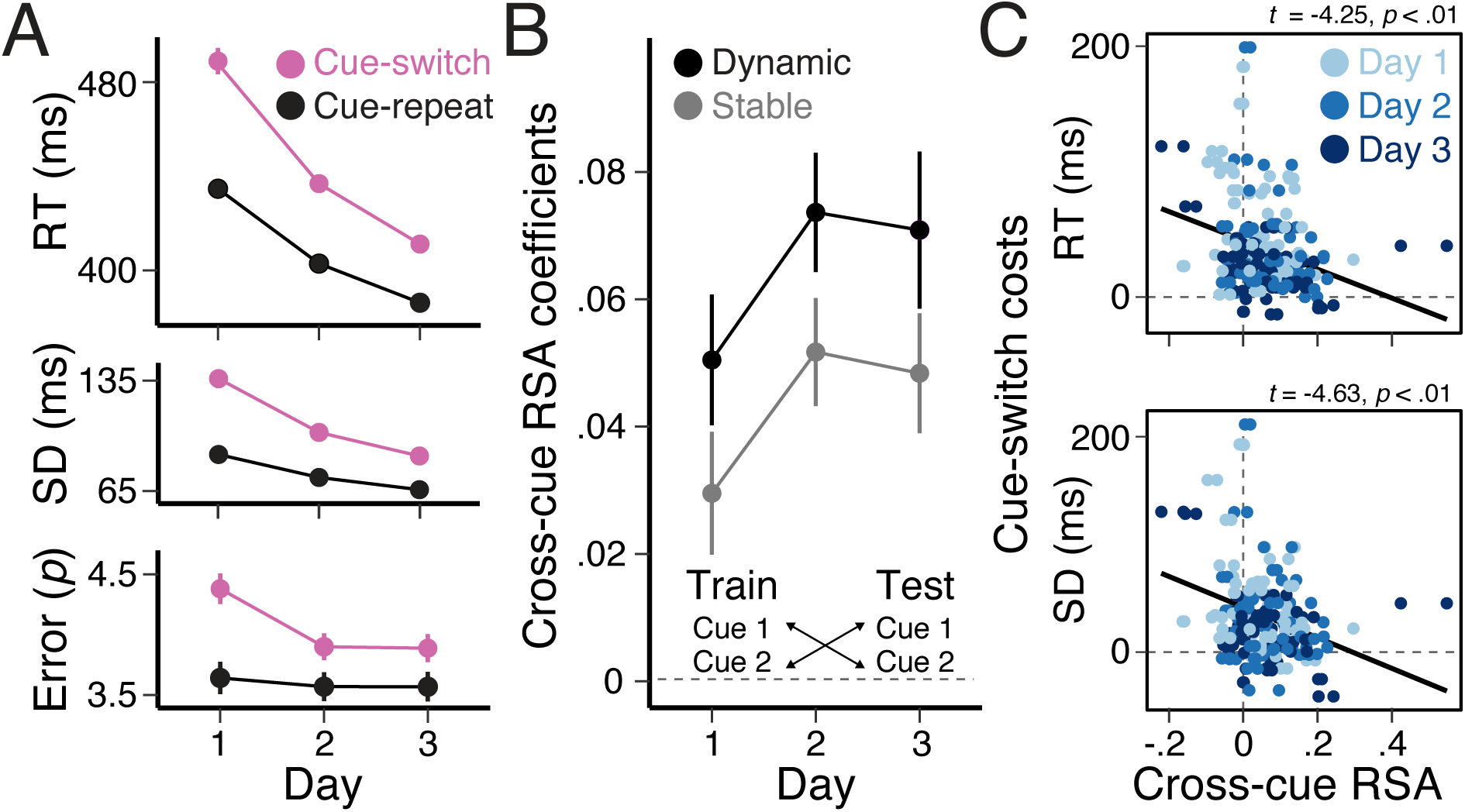
Cue-switching costs and cross-cue generalization. (A) Costs of switching cues in trials with the repetitions of task rules, stimulus, and response. (B) The changes in average RSA scores of rule-specific conjunctions that generalize over cues across days. The temporal generalization RSA is performed in a cue-specific manner, and then the results are averaged over 0-300ms time periods. (C) Correlations between behavioral costs of switching cues (i.e., the average and variability of RTs) and the cross-cue generalization decoding of rule-specific conjunctive subspaces.

Further, these changes were behaviorally relevant. In participants showing more generalized patterns of context-specific conjunctions, the costs of RTs and the variability of RTs tended to be lower (Figure 6C; see Supplementary Figure 3 for other task features; RT cue-switch costs: *t*(1,38) = -4.26, *beta* = -.28, 95 % CI [-.41 -.15]; SD cue-switch costs: *t*(1,38) = -4.62, *beta* = -.30, 95 % CI [-.43 -.17]). Further, such differences among individuals become progressively stronger across days (RT cue-switch costs: *t*(1,38) = -2.87, *beta* = -.39, 95 % CI [-.66 -.12] for linear effect; *t*(1,38) = - 2.07, *beta* = -.26, 95 % CI [-.49 -.02] for quadratic effect; SD cue-switch costs: *t*(1,38) = -4.62, *beta* = -.30, 95 % CI [-.50 -.01] for linear effect; *t*(1,38) = -1.74, *beta* = -.20, 95 % CI [-.43 .26] for linear effect). Taken together, these results indicate that extensive practice reshaped the neural dynamics encoding context-specific conjunctions to be more stable and abstract, providing more efficient access to the subspace corresponding to the minimally sufficient task-tailored solutions.

## Discussion

What changes in neural task representations lead to performance gains over the course of extensive practice? We report two results that give novel insights into this question. First, we found that accumulated experience with specific task instances over the course of a session – defined in terms of a particular co-occurrence of stimulus, response, and rule context – led to the strengthening of high-level context-specific conjunctive representations. Over the course of practice, the dynamics encoding these representations became stabilized at progressively earlier points in time with respect to the beginning of trials, and they contributed to increased dimensionality of task representations. The strengthening of these conjunctive representations, and not other levels of representation, accounted for the power law-like improvement in performance during early phase of learning. Second, we observed that while conjunctions were strengthened, they were also shaped over days so that dynamics became generalizable (i.e., aligned) over redundant task cues. These changes were correlated with reductions in the costs of switching task cues. Taken together, our results indicate that practice optimizes stable encoding and retrieval of task-tailored context-specific conjunctive representations. We now consider each finding in turn.

On each trial of the task, participants encountered a specific combination of rule, stimulus, and response. As experience with these trial events accumulated over the course of a session, we observed that non-linear conjunctive representations of the combination of these features increased in strength with each encounter (Figure 3B). We also saw a similar increase in strength of the stimulus and response representations with experience, but critically not for S-R conjunctions. However, only the changes in context-specific conjunctions accounted for the power law-like rapid improvement in performance in the first session.

Different theories of skill learning, habits, and automatic behavior have emphasized learning at different levels of task representation. A long tradition in the study of cognitive control, for instance, has emphasized the learning of context-independent stimulus-to-response (S-R) mappings as the basis of automatic behavior (Miller & Cohen, 2001). Information about rules and contexts maintained in working memory can bias the activation of task-relevant S-R associations over competitors, which serves as the basis for controlled behavior. Similarly, several models of motor learning assume learning of S-R mappings is at the base of practice-based change to gain efficiency (Du et al., 2022; Hardwick et al., 2019; Wood & Rünger, 2016).

Similarly, chunking theories of automaticity assume that the benefit of learning occurs first in more general and then more specific combinations or chunks (A. Newell, 1993). This would predict that learning of context-independent S-R conjunctions (simpler chunks) would drive performance change early in the session, while learning of context-specific conjunctions (higher-order chunks) would affect improvement late.

However, we did not see changes in the S-R conjunctions that related to performance gains at any stage. This was unlikely due to our task design. Rather, our task was designed so that particular S-R mappings were encountered across different contexts. Therefore, any S-R conjunctions were experienced twice as frequently as context-specific conjunctions, offering ample opportunity for experience-based improvement. Nevertheless, we did not observe a systematic increase in the strength of these S-R conjunctions over the course of practice, nor did their strength relate to behavioral change.

In this sense, the present observations align most closely with the assumptions of instance theory (Gordon D. Logan, 1988) that assigns learning to high-level, specific task events. According to this theory, a task can be performed initially by way of a slow but flexible algorithm that can assemble any task. However, each time a task is performed, a memory of the specific combination of task features during each processing event is obligatorily encoded as an instance in memory. On future attempts to perform the task, the slow algorithm races with the time it takes to retrieve an instance. As more instances accrue, random sampling of faster retrieval is more likely to occur, winning the race (G. D. Logan, 1992). However, given a distribution of retrieval times, retrieval events that are more extreme become rarer over time, resulting in a decelerating performance improvement toward an asymptote.

In line with these ideas, our results relate practice-based improvement only to facilitated access to high-dimensional, context-specific conjunctions. Indeed, we observed changes in the neural subspaces that we have previously associated with facilitated context-dependent readout (Badre et al., 2021; Kikumoto & Mayr, 2020; Kikumoto, Mayr, et al., 2022; Kikumoto, Sameshima, et al., 2022; Rangel et al., 2023). Specifically, in prior work (Kikumoto et al., in Press), we used a response-deadline procedure to characterize the representational state that facilitated correctly responding in the virtually same task used here. Using this procedure, we found that fast and accurate task performance followed from the availability of conjunctive representations that were encoded within a high-dimensional neural geometry and that had transitioned from a temporally dynamic to a stable state.

In the present study, we found these same features improved with practice. Specifically, a stronger conjunctive representation was encoded within a higher dimensional geometry over sessions, and this representation progressively stabilized at earlier timepoints relative to the response. Thus, representational changes that facilitate access to the conjunctive representation to guide efficient performance are directly enhanced with repeated practice.

Over the course of days, we also observed that the conjunctive representation changed to be generalizable over task features that are salient but redundant for defining response-relevant contingencies in the task. In particular, in this task, we included two cues for each rule. For example, the vertical rule was cued by both the words “vertical” and “updown”. Early in practice, relatively independent geometries encoded conjunctions for different rule cues, even though the rule context and S-R mapping (i.e., context-specific conjunction) were shared. Switching these redundant task cues incurred significant costs on behavior. However, over the course of practice, these distinct conjunctive subspaces came to align, forming one conjunctive rule subspace that abstracted over differences in the redundant rule cues. This abstraction predicted a gradual reduction of behavioral cost from switching the cue.

Accordingly, in the current study, we show that high-dimensional representations that guide efficient responding are not always incidental bindings of all task-relevant features at a particular moment in time, but rather, come to represent neural states that are shaped based on abstract task demands. They incorporate only the task-relevant features needed to minimally separate task contingencies to make a response, and slowly, form abstractions over unnecessary redundant elements. However, unlike the changes in the strength of context-specific conjunctive code, these changes to its representational geometry occurred over the course of sessions, rather than within a session. We return to this distinction below.

It is notable that the abstract alignment of context-specific conjunctions is inconsistent with the strictest version of instance theory (Logan, 1988). In its original formulation, instances were akin to episodic memory-like bindings of all features of a task event (e.g., task cues or timing of trials). Nonetheless, there is neither a mechanism for transforming these instances based on redundant features of the input, nor a clear prediction for how the system determines the complexity of instances given the tasks. Similarly, this kind of abstraction is not a feature of chunking or other theories of automaticity. However, these theories mostly focused on improvement in power law which occurred only in session 1 in our study, and so in some sense did not need this kind of abstraction for their explanatory scope. Thus, this observation raises the question—why does this abstraction occur?

There are several reasons abstraction of this kind would be advantageous. First, abstraction is a basis for generalization which can provide robustness against interference (e.g., input variability) and can allow reuse of representations for new task scenarios or transfer to new tasks. Second, coding extra dimensions may be costly in several ways, and so abstraction may be encouraged by some form of regularization (Wojcik et al., 2023). Finally, abstraction allows separate instances of action contexts to be treated as one, allowing them to benefit from repetition, which leads to performance benefits. Indeed, our data provide evidence of this last advantage, showing that abstraction over the rule cue predicted reduced cue switch costs over sessions.

Collectively, then, these results provide an answer to how task representational geometry changes with practice, and specifically to the question of whether dimensionality expands or reduces over the course of practice. Paradoxically, both kinds of change occur. There is progress toward higher-dimensional geometry driven by an enhanced access to task-tailored (i.e., structured not random), context-specific conjunctions. These changes reflect greater separation among conjunctive subspaces, i.e., higher dimensionality, as well as changes in features like temporal stability.

However, there is also shaping away from diverse higher dimensional via heterogenous subspaces toward a more efficient and abstract (i.e. lower-dimensional) space of conjunctions within the structure and objective of task contingencies.

Thus, rather than conceptualizing automaticity as progress toward higher or lower dimensional task representations. Our results suggest that practice optimizes access to a tailored task representation that incorporates combinations of all those features, and only those features, needed to specify a correct output to solve each instance of the task at hand. Thus, in time, this accessible, high-dimensional code features a library of specific task events that tessellate the full task space (as defined by the task’s objective function), but not necessarily the full input space.

A key open question concerns what mechanisms of learning underlie the changes we observe and why. The pattern of optimization we observe here is in line with what would characterize representations found in neural network models trained to solve a context-dependent response selection task (Farrell et al., 2022). If learning abstract dynamics reduces large costs in task performance (Figure 6), then gradient-based optimization should discover such solution manifolds (Driscoll et al., 2024; Goudar et al., 2023; Sandbrink & Summerfield, 2024; Saxe et al., 2022). However, gradient-based methods (e.g., backpropagation) and their approximations may not accurately represent what occurs during extensive practice, nor are they necessarily desirable (Scott & Frank, 2021). Nevertheless, plausible biological mechanisms, such as three-factor plasticity combined with noise correlation (Frémaux et al., 2013; Nassar et al., 2021), are known to support comparable optimizations that promote robustness against interference and enhance generalization.

Regardless of the types of algorithms, however, a further open issue is that optimization requires an objective function to be defined and maintained without interference from other objectives and shared within the system. While this is straightforward to implement in artificial agents within well-controlled learning contexts, it is not clear how such an objective of learning is implemented in a human, even when practicing one task (e.g., an instructed experimental task), alongside many other tasks and objectives in various contexts (e.g., doing homework). Thus, our data establish that extensive practice may be understood as a process of optimizing access to task-tailored conjunctive codes across conditions of a task, but it is not clear how practice yields this change.

While our data cannot establish the mechanisms of learning, our results do offer potential clues as a basis for future research. In particular, we observed a key distinction in the time course of practice-based change between access to the conjunction versus shaping of the conjunction. Again, changes in the strength and accessibility of the conjunctive representation only explained the within-session power law-like improvement in performance. We did not see a relationship between these changes and the offline gains in performance observed for each session. Conversely, abstraction of the conjunctive subspace over the redundant task cues input unfolded over multiple sessions that were separated by long rest periods including a night of sleep. That suggests that reshaping of the dynamic geometry of task representations occurs at multiple temporal and representational scales (Kikumoto & Mayr, 2018) and might utilize different mechanisms.

While abrupt, non-linear learning (e.g., after rest) is ubiquitously observed in visual perceptual learning and motor learning tasks (Censor et al., 2012; Tamaki et al., 2020; Walker et al., 2002), we did not predict this difference in time course, and so it will be important to replicate this observation. Nevertheless, this difference might point at an important difference in learning. For example, changes in separation and stability that facilitate access to conjunctive representations might be driven by performance-dependent mechanisms like Hebbian plasticity in cortical synapses or episodic encoding mechanisms through the hippocampus that occur within the session (Mill & Cole, 2023). In contrast, the changes leading to the abstraction over irrelevant features might require latent consolidation mechanisms, such as those associated with the formation of abstract schemas in memory (Behrens et al., 2018; Preston & Eichenbaum, 2013; Rasch & Born, 2013), statistical regularizations of the learned weights (Löwe et al., 2024), or replay (Preston & Eichenbaum, 2013) and synaptic downscaling (Norimoto et al., 2018) that are often regulated by over night of sleep. Future work manipulating the conditions of practice or overnight consolidation might provide insights into these mechanisms of change.

There are limitations in the present study that put our results in context. A first consideration is the generalizability of the present conclusions. We tested only one task, and as such, we leave open the possibility that other tasks would lead to different outcomes. It will be important to replicate the present results in other tasks and paradigms, particularly in settings that are more complex, temporally extended, demanding of working memory, and/or ecologically valid. Nevertheless, the present task is designed to test how different responses are selected to the same stimulus dependent on a contextual rule. That context-sensitivity sets this task apart from the S-R learning tasks commonly used in the motor learning literature (Hardwick et al., 2019). And, this basic context-dependent rule structure is representative of most, if not all, tasks requiring cognitive control at their most reduced level.

Further, while the task lends itself to focusing on context-dependent conjunctions, we also included the opportunity to learn more low-dimensional intermediate representations, like the common S-R mappings, that were shared over more trials and so could have facilitated behavior under some models of practice-related improvement. But, again, although S-R conjunctions were correlated with trial-to-trial performance, they were not consistently associated with practice-related change. Thus, we believe the current observations will generalize to many other tasks and reflect general properties of practice-related change in goal-directed behavior requiring context-sensitivity.

While we view our observations to be generalizable to other cases of practice of a stable, context-dependent rule structure, we have not tested all aspects of practice and skill learning, and so we have not identified the only drivers of practice-related change in performance. For example, changes in task rules and strategies (e.g., discovering that going over the bar backward improves the high jump) will result in abrupt shifts in task performance (Schuck et al., 2015; Taylor & Ivry, 2012). Such changes can be discovered through experience, and our results do not address such effects. Rather, we view these observations as relating to refinement and optimization given a particular stable strategy, objective and structure of the task, thereby expressing how target representational states change to facilitate their access and match to task contingencies over the course of practice.

In summary, we observe that experience with a task enhances access to context-integrated conjunctive representations that correspond to specific combinations of rule, stimulus, and response during each task event within a task space, as defined by a task’s objective function. This facilitated access accounted for the power law-like improvements observed within a session of practice. Over the course of multiple sessions, this target conjunctive representation was shaped to be encoded in temporally stable dynamics and abstract neural subspaces. These results provide a new window on the targets of practice-related change that leads to performance improvement on a task.

## Acknowledgments

We would like to acknowledge the members of the Human Cognition and Learning lab at RIKEN, particularly, Honma Saki, Sara Matsui, Minori Fukushima, and Norie Kawamura. Further, we would like to acknowledge the members of the Badre lab, the laboratory of neural computation and cognition, Learning Memory & Decision lab and the Shenhav lab. This project was supported by funding from the National Institute of Mental Health (R01 MH125497), the National Institute of Neurological Disorders and Stroke (R21 NS108380), and a Multidisciplinary University Research Initiative award from the Office of Naval Research (N00014-16-1-2832) to DB and from JSPS KAKENHI Grant Number 19H01041, 20H05715, JST Moonshot R&D JPMJMS2013 to KS and the JSPS Overseas Research Fellowships to AK.

## Author Contributions

Project conception: A.K and D.B; Methodology: A.K; Data Collection: A.K, T.N, and K.S.; Analysis and Interpretation: A.K and D.B.; Manuscript writing: A.K, K.S, and D.B; Funding acquisition: K.S and D.B; Supervision: K.S and D.B.

## Supplementary Figures

**Figure S1.**
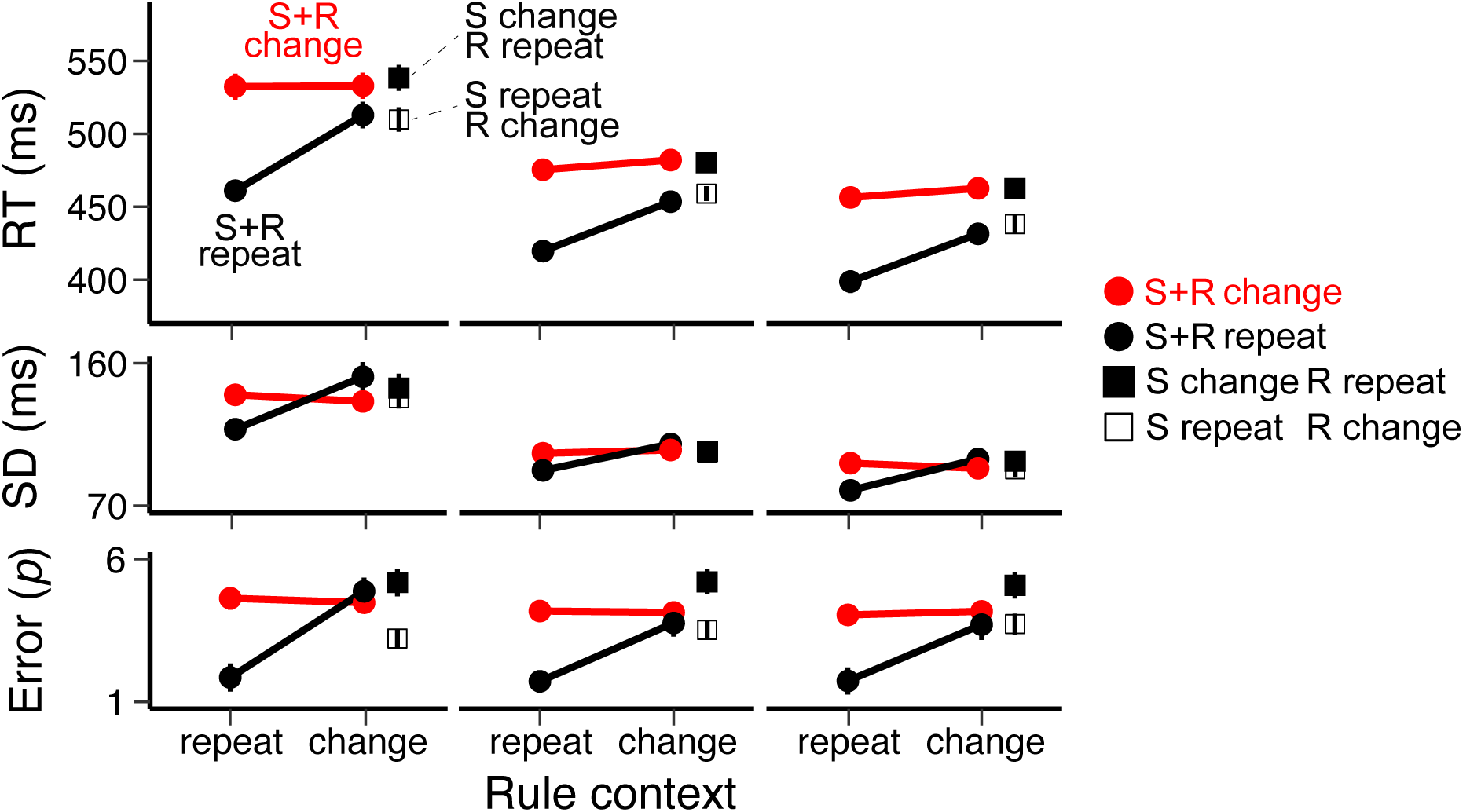
Partial overlap costs across sessions. Average RTs, variability of RT, and errors as a function of changes in rule, stimulus, and response features.

**Figure S2.**
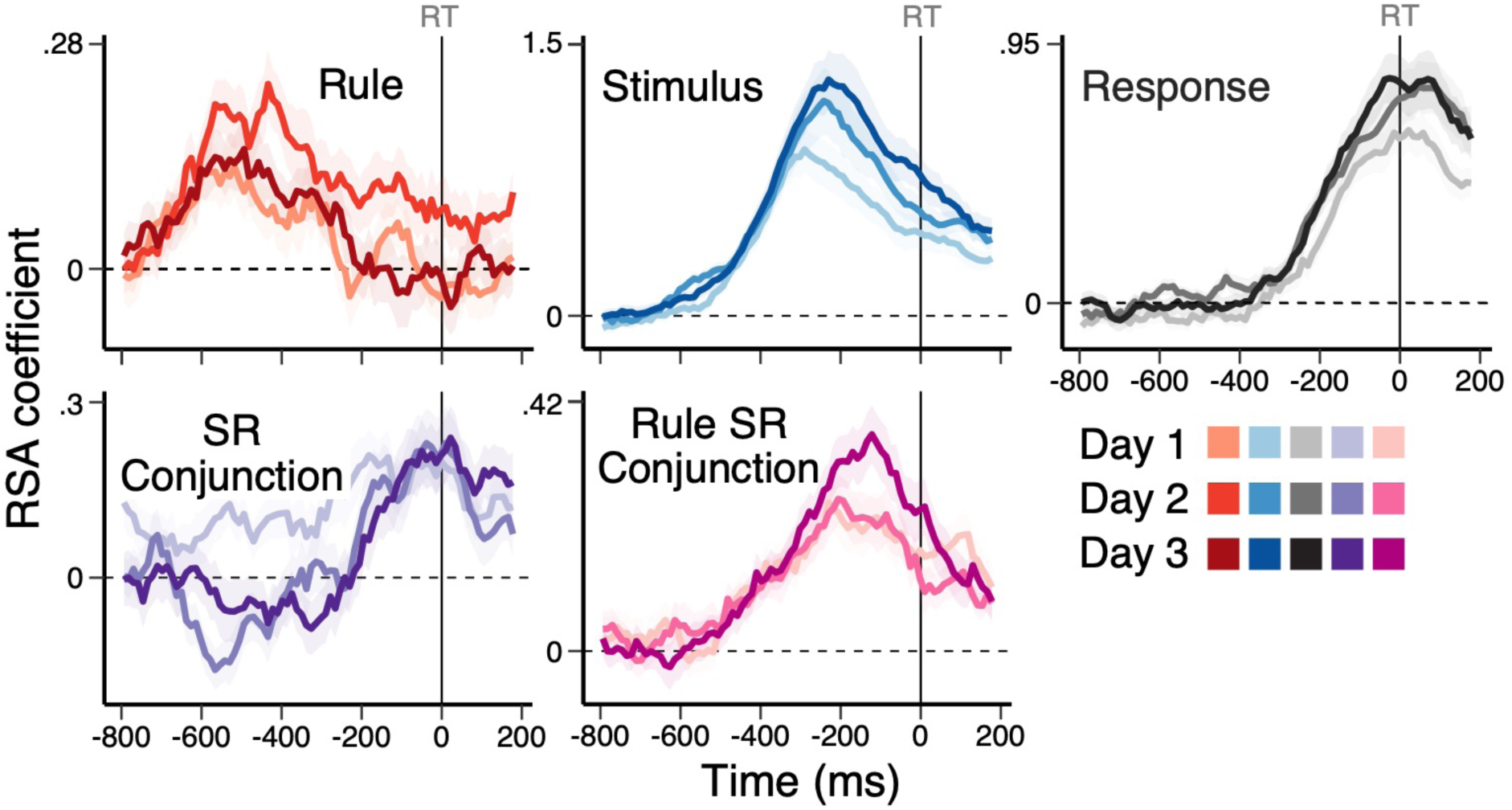
Response-aligned time-course of decoding of task representations. Average, single-trial RSA coefficients (*t*-values) associated with each of the basis set task features (rule, stimulus, and response) and two conjunctions (SR conjunction and rule-specific SR conjunction) that are aligned to the onset of the response execution.

**Figure S3.**
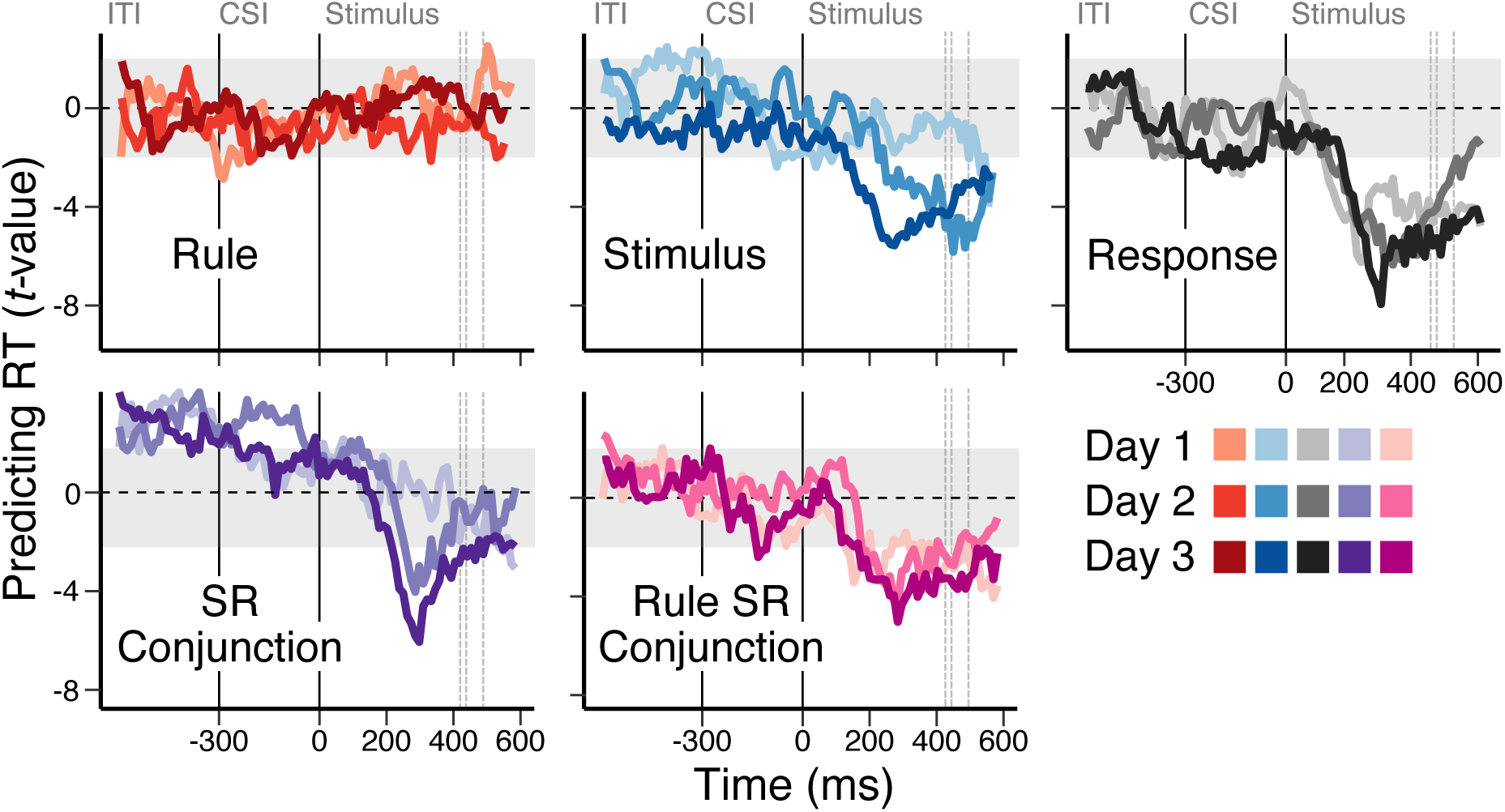
Time-resolved regression of RTs by decoded task representations. The correlation between single-trial RT and RSA coefficients of rule, stimulus, response, SR conjunction, and rule-specific SR conjunction. A series of multi-level models were fitted within each session separately, using all RSA coefficients simultaneously.

**Figure S4.**
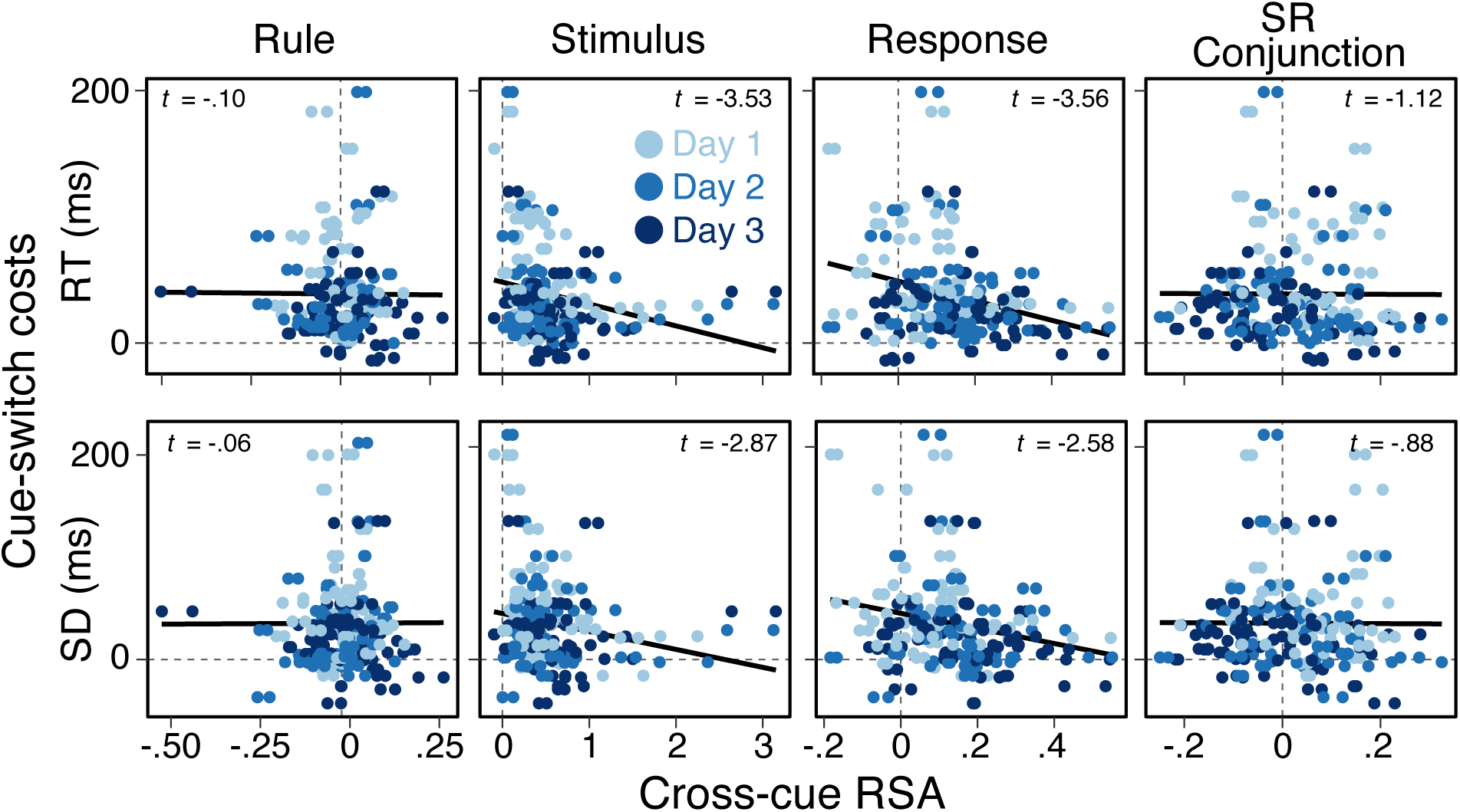
Cue-switching costs and cross-cue generalization. Correlations between behavioral costs of switching cues (i.e., the average and variability of RTs) and the cross-cue generalization decoding of rule, stimulus, response and stimulus-response subspaces.

